# Diffusion-mediated quantification of dose-dependent antifungal drug tolerance

**DOI:** 10.1101/2025.09.05.674536

**Authors:** Samira Rasouli Koohi, Clare Maristela Galon, Daniel A. Charlebois

## Abstract

Antimicrobial resistance is a global health problem, with drug-resistant fungi posing major challenges for the treatment of immunocompromised patients. Antifungal tolerance is a recently discovered phenomenon whereby pathogenic fungi, including the multidrug-resistant pathogenic yeast *Candidozyma auris* (formerly *Candida auris*), grow slowly above minimum inhibitory drug concentrations. We combine physics-based spatiotemporal models with microbiology experiments to quantitatively investigate the emergence of tolerance to all three major classes of antifungal drugs in *C. auris*. Specifically, we combine Fick’s second law solved using a finite difference method to simulate drug diffusion with data from experimental disk diffusion assays to determine the concentration ranges at which antifungal-tolerant colonies emerge. This biophysics study advances antimicrobial resistance research by providing a method to quantify antifungal tolerance and by demonstrating that antifungal tolerance is a strain-, drug-, and dose-dependent phenomenon in an emerging fungal pathogen.

## 1. INTRODUCTION

Antimicrobial resistance (AMR) is a global health problem with 1.27 million deaths attributable to AMR in 2019 [1]. Fungal pathogens resistant to antifungal drugs substantially contribute to AMR [2]. There are unique challenges associated with the treatment of fungi compared to other microbes, including a limited number of classes of antifungal drugs (azoles, polyenes, and echinocandins) approved to treat invasive infections in humans [3, 4] and thermal adaptation and range expansion due to climate change [5]. Specific challenges in antifungal drug development include the similarity between eukaryotic fungal cells and human cells, which restricts drug selectivity and increases toxicity [6], as well as the ability of pathogenic fungi to form biofilms that reduce drug penetration [7]. Despite the fact that fungal diseases cause 1.7 million deaths annually [8], research on antifungal resistance has lagged behind research on antibacterial resistance [9].

Particularly concerning is the emergence of multidrug and pan-resistant yeast pathogens, especially *Candida auris* (recently reclassified as *Candidozyma auris* [10]) [11–14]. Candidemia, a type of bloodstream infection caused by *C. auris* and related *Candida species*, can cause organ damage [15]; the hospital mortality rate for candidemia is estimated to range from 30 % to 70 % [16, 17]. Antifungal resistance in *C. auris* poses treatment challenges, with high rates of resistance observed against azole, polyene, and echinocandin antifungal drugs [18, 19]. Accordingly, the WHO listed *C. auris* as a critical priority fungal pathogen in 2022 [20]. Mechanisms of antifungal resistance in *C. auris* include genetic mutations affecting drug targets (e.g., ERG11), efflux pumps (e.g., MDR1), and alterations in cell wall composition [21, 22].

Each antifungal drug belongs to a distinct class with a unique mechanism of action [23, 24]. For instance, itraconazole, an azole antifungal drug, inhibits the enzyme lanosterol 14*α*-demethylase required for the biosynthesis of ergosterol. This inhibition disrupts the integrity of the cell membrane and suppresses fungal cell growth and reproduction (fungistatic drug activity). Amphotericin B, a member of the polyene drug class, promotes fungal cell death (fungicidal drug activity) by binding to ergosterol, leading to pore formation and loss of membrane integrity. Caspofungin, a member of the echinocandin drug class, targets the enzyme *β*-1,3-glucan synthase, a core component of the fungal cell wall. Caspofungin exhibits fungicidal activity against *C. auris* and most *Candida* species [23].

Antifungal tolerance is a recently discovered phenomenon, whereby fungi grow slowly above the minimum inhibitory concentration (MIC) of an antifungal drug [25, 26]. Tolerance can be quantified experimentally using either a broth microdilution assay (BMDA) for liquid cultures or a disk diffusion assay (DDA) for semi-solid agar cultures [27]. Tolerance was shown to predict fluconazole efficacy in patients with *Candida albicans* bloodstream infections [28]. Despite this, we lack clinical diagnostic tests to detect tolerance [29, 30]. Antifungal tolerance is distinct from antifungal resistance [26] in that 1) Resistance is a permanent/heritable in which MIC changes, whereas tolerance is a reversible phenomenon in which MIC does not change [27, 30]; 2) Resistance is associated with specific genetic changes that affect a limited number of known pathways, whereas tolerance is thought to be governed by general stress responses and affected by many pathways [29, 31, 32]; and 3) Resistant cells grow quickly above MIC, have short lag times in drug conditions [33], and colonies appear within 24 hours in DDAs, whereas tolerant cells grow slowly above MIC, have more variable lag times in drug conditions, and emerge around 48 hours in DDAs [27, 30].

Previous experimental research on the same panel of clinical *C. auris* isolates combined BMDAs and DDAs with image analysis to quantify tolerance as supra-MIC growth (SMG) and fraction of growth (FoG) within the zone of inhibition (ZOI) [30]. This work established that 1) *C. auris* is tolerant to fungistatic azole and fungicidal polyene and echinocandin antifungal drugs; 2) The drug-tolerant *C. auris* phenotype is reversible to the drug-susceptible phenotype; and 3) Tolerance and resistance in some *C. auris* isolate–antifungal combinations can be reduced or eliminated by combining antifungal drugs with the adjuvant chloroquine. These results highlight the biological and clinical relevance of antifungal tolerance and motivate the need for a quantitative framework that relates drug concentration to the emergence of tolerant subpopulations. Another experimental study on *Candida albicans* found that high concentrations of fluconazole select for antifungal tolerance, whereas low fluconazole concentrations select for antifungal resistance [25]. This suggests that tolerance, a phenomenon that is currently overlooked in clinical antifungal susceptibility testing [27, 30], can emerge as a dominant adaptive strategy under high selective pressure [25]. Despite the advances made by this research, the dose-dependent nature of antifungal drug tolerance in pathogenic yeast species other than *C. albicans* has not been investigated.

On the computational side, a finite difference computational model based on Fick’s second law of diffusion was used to predict the radius of the ZOI (RAD) in diffusion-based bioassays [34]. Subsequently, the MIC based on RAD measurements was calculated without prior knowledge of the diffusion rate [32]. However, drug diffusion modeling has not been used to quantify the drug concentrations at which tolerance emerges.

In this study, we address the above knowledge gaps by combining physics-based spatiotemporal diffusion models with microbiological DDA experiments to quantitatively investigate the emergence of tolerance in the pathogenic yeast *C. auris* to all three major classes of antifungal drugs. First, we use finite difference method simulations based on Fick’s second law to predict the diffusion-mediated spatiotemporal concentration profiles of the antifungal drugs itraconazole, amphotericin B, and caspofungin on semi-solid agar. Then we combine these simulated antifungal drug concentration profiles with our experimental DDA data to determine the drug concentration ranges at which antifungal tolerant *C. auris* colonies emerge. Overall, we develop a diffusion-based method to quantify the drug concentration ranges at which antifungal tolerance emerges *in vitro* and find that antifungal tolerance is a strain-, drug-, and dose-dependent phenomenon in an emerging human fungal pathogen.

## II. MODELING & SIMULATION

We used quantitative modeling and simulation to investigate how diffusion-mediated spatiotemporal drug concentrations affect the emergence of tolerant *C. auris* colonies in DDAs.

### A. Drug diffusion model

As yeast colonies are constrained to grow on the surface of semi-solid agar, we began by modeling antifungal drug diffusion using the two-dimensional diffusion equation [35]:

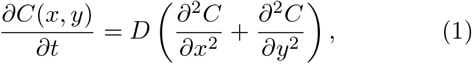

where *C* is the antifungal drug concentration and *D* is the diffusion constant.

Here, we assumed pure diffusion of drug molecules and did not explicitly consider any other mechanisms that can influence the drug concentration profile, such as degradation, active transport, advection, or evaporation (see Sec. II B) [36, 37]. This assumption has been shown to be valid for the diffusion of antifungal drugs in agar [32, 38– 40]. Active transport, advection, and medium-mediated degradation are negligible in DDAs as agar is a stable, semi-solid medium in which chemical degradation and active fluid flow are absent. As DDAs are performed using a closed Petri dish in a humidified incubator, evaporation is minimal. These factors support diffusion as the dominant mechanism of mass transfer, producing a reasonable model of drug diffusion dynamics in DDAs [32, 38–40].

To determine an initial seed value of the diffusion constant (*D*^*′*^) for our simulations (see Sec. II B), we used the Stokes-Einstein relation:

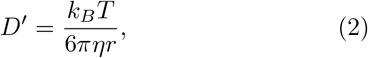

where *k*_*B*_ is the Boltzmann constant, *T* is the temperature, *η* is the viscosity of the agar medium, and *r* is the radius of a spherical drug molecule. The molecular weights of itraconazole (*m*_*itr*_), amphotericin B (*m*_*amp*_), and caspofungin (*m*_*cas*_), were used to calculate the radii of these drug molecules (*r*_*itr*_, *r*_*amp*_, and *r*_*cas*_; Table II in Appendix A) [41]:

**TABLE I.** Antifungal drug concentration ranges and radii at which tolerant colonies emerged for different *C. auris* isolates and antifungal drugs. *C*_1_ and *C*_2_ are the simulated drug concentrations corresponding to the experimental radii *R*_1_ and *R*_2_ of tolerant colonies closest and furthest from the antifungal disk, respectively, after 48 hours.

| ID | Isolate No. | Antifungal Drug | $C_1 - C_2$ ($\mu\text{g/ml}$) | $R_1 - R_2$ (mm) | $C$ Change (%) | $R$ Change (%) ± Error |
| --- | --- | --- | --- | --- | --- | --- |
| ID1 | 1 | Caspofungin | 0.641 - 0.415 | $8.0 \pm 0.5 - 12.5 \pm 0.5$ | 35.3 | $56.3 \pm 11.6$ |
| ID2 | 2 | Caspofungin | 0.751 - 0.415 | $5.5 \pm 0.5 - 12.5 \pm 0.5$ | 44.7 | $127.3 \pm 24.9$ |
| ID3 | 1 | Amphotericin B | 0.979 - 0.922 | $9.0 \pm 0.5 - 11.0 \pm 0.5$ | 5.82 | $22.2 \pm 8.3$ |
| ID4 | 1 | Itraconazole | 4.07 - 2.74 | $5.5 \pm 0.5 - 14.0 \pm 0.5$ | 32.7 | $154.6 \pm 29.0$ |

**TABLE II.**
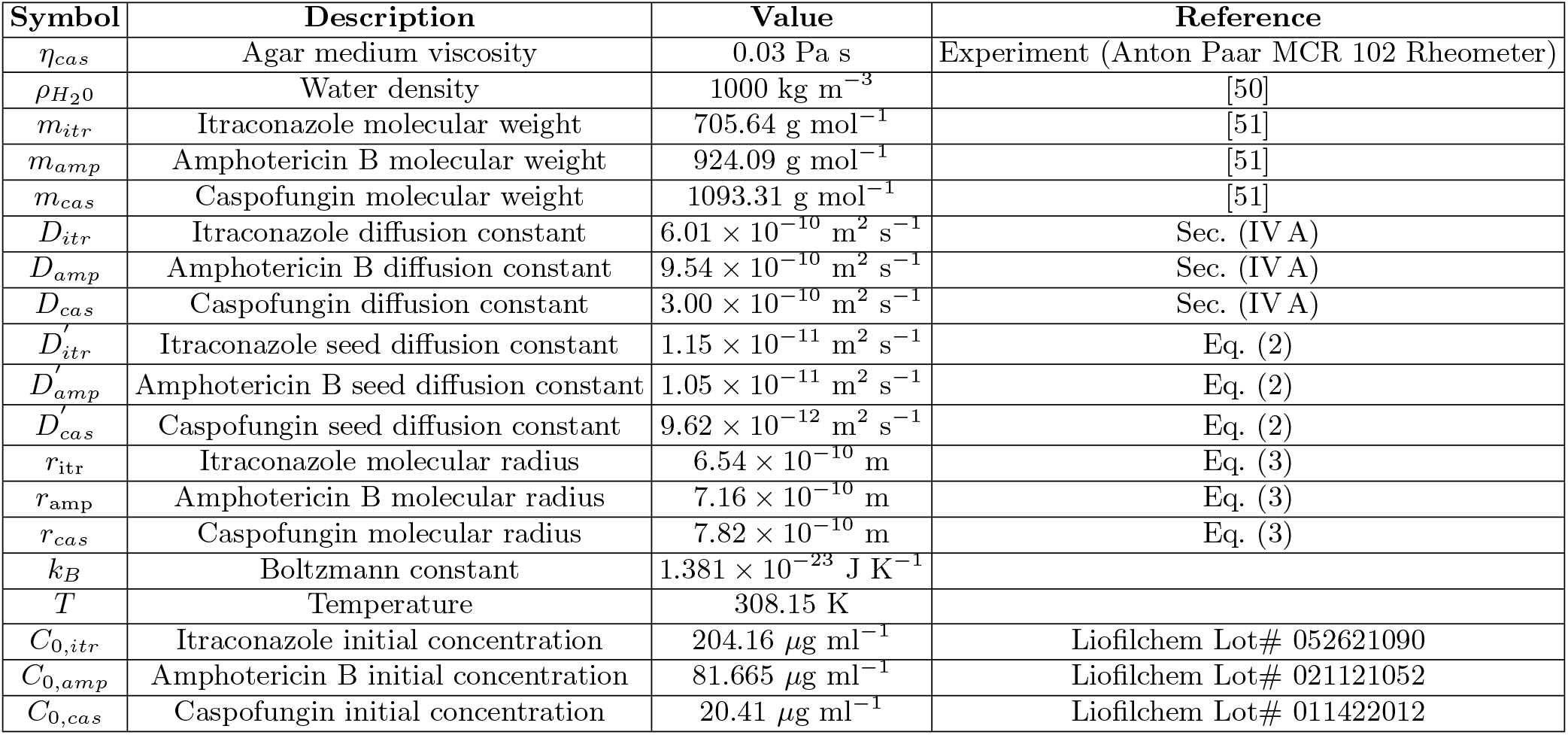
Parameters for modeling and numerically simulating the diffusion of caspofungin, itraconazole, and amphotericin B in agar medium. 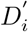 denotes an initial, calculated diffusion constant for an antifungal drug *i. D*_*i*_ a represents a more accurate, computationally determined diffusion constant for an antifungal drug *i*.

| Symbol | Description | Value | Reference |
| --- | --- | --- | --- |
| $\eta_{cas}$ | Agar medium viscosity | 0.03 Pa s | Experiment (Anton Paar MCR 102 Rheometer) |
| $\rho_{H_2O}$ | Water density | 1000 kg m<sup>-3</sup> | [50] |
| $m_{itr}$ | Itraconazole molecular weight | 705.64 g mol<sup>-1</sup> | [51] |
| $m_{amp}$ | Amphotericin B molecular weight | 924.09 g mol<sup>-1</sup> | [51] |
| $m_{cas}$ | Caspofungin molecular weight | 1093.31 g mol<sup>-1</sup> | [51] |
| $D_{itr}$ | Itraconazole diffusion constant | $6.01 \times 10^{-10}$ m<sup>2</sup> s<sup>-1</sup> | Sec. (IV A) |
| $D_{amp}$ | Amphotericin B diffusion constant | $9.54 \times 10^{-10}$ m<sup>2</sup> s<sup>-1</sup> | Sec. (IV A) |
| $D_{cas}$ | Caspofungin diffusion constant | $3.00 \times 10^{-10}$ m<sup>2</sup> s<sup>-1</sup> | Sec. (IV A) |
| $D'_{itr}$ | Itraconazole seed diffusion constant | $1.15 \times 10^{-11}$ m<sup>2</sup> s<sup>-1</sup> | Eq. (2) |
| $D'_{amp}$ | Amphotericin B seed diffusion constant | $1.05 \times 10^{-11}$ m<sup>2</sup> s<sup>-1</sup> | Eq. (2) |
| $D'_{cas}$ | Caspofungin seed diffusion constant | $9.62 \times 10^{-12}$ m<sup>2</sup> s<sup>-1</sup> | Eq. (2) |
| $r_{itr}$ | Itraconazole molecular radius | $6.54 \times 10^{-10}$ m | Eq. (3) |
| $r_{amp}$ | Amphotericin B molecular radius | $7.16 \times 10^{-10}$ m | Eq. (3) |
| $r_{cas}$ | Caspofungin molecular radius | $7.82 \times 10^{-10}$ m | Eq. (3) |
| $k_B$ | Boltzmann constant | $1.381 \times 10^{-23}$ J K<sup>-1</sup> | |
| $T$ | Temperature | 308.15 K | |
| $C_{0,itr}$ | Itraconazole initial concentration | 204.16 μg ml<sup>-1</sup> | Liofilchem Lot# 052621090 |
| $C_{0,amp}$ | Amphotericin B initial concentration | 81.665 μg ml<sup>-1</sup> | Liofilchem Lot# 021121052 |
| $C_{0,cas}$ | Caspofungin initial concentration | 20.41 μg ml<sup>-1</sup> | Liofilchem Lot# 011422012 |

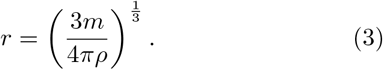

The density of each antifungal drug *ρ* was assumed to have the same density as water 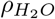. This assumption is based on the fact that each antifungal drug considered was highly diluted in water to obtain the final drug concentration for our DDA experiments (Sec. III).

### B. Numerically simulating drug diffusion

We used a finite difference method [42] to numerically solve the diffusion equation [Eq. (1)], which was used to model drug diffusion across the agar surface. An agar plate was represented by a grid of points (*i,j*), each with associated spacing Δ*x* and Δ*y* in the *x* and *y* directions, along with time spacing Δ*t*, to approximate the drug concentration *C*(*x, y, t*):

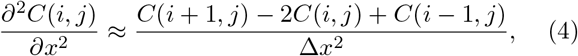

and

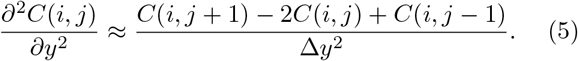

Substituting Eqs. (4) and (5) into Eq. (1) yields a finite difference equation:

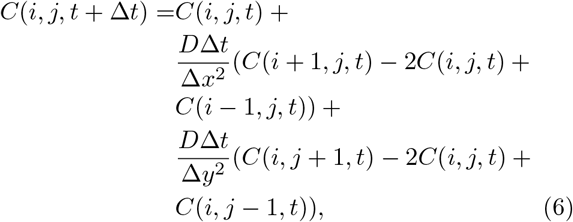

which relates *C*(*i, j, t* + Δ*t*) to *C*(*i, j, t*). The Petri dish was modeled as a square with dimensions 85 mm x 85 mm with Δ*x* = Δ*y* = 0.1 mm.

The numerical simulations were initialized by setting the antifungal drug concentration at the center of the grid to the initial concentration of the drug disk (*C*_0,*itr*_, *C*_0,*amp*_, and *C*_0,*cas*_; Table II in Appendix A) used in the DDA experiments (Sec. III). The drug concentration at all other grid points was initially set to zero. At each simulation time step, the drug concentration at each grid point was calculated using the drug concentrations of the neighboring grid points from the previous time step [Eq. (6)]. This numerical approach generated time-dependent antifungal drug concentration profiles from which we determined the drug concentrations at which drug-tolerant colonies emerged in our DDA experiments (Sec. IV).

To determine a value of *D* that corresponded to our DDA experiments, we assumed that the minimum fungicidal concentration (MFC; lowest concentration of an antifungal drug that kills susceptible fungal cells) corresponded to the drug concentration at the edge of the ZOI after 48 hours for each *C. auris* isolate-antifungal drug combination. The edge of the ZOI in a DDA represents the boundary beyond which antimicrobial drugs no longer negatively affect susceptible cells. To facilitate comparison between our simulated surface area-based drug concentrations and our experimentally determined volumetric-based MFCs, we divided the simulated 2D drug concentrations by the depth of the agar (8.66 mm). Simulations using the theoretically determined diffusion constant (Eq. 2) did not produce drug concentration profiles that agreed with our experimental ZOI-MFC observations (Sec. IV A). Therefore, we iteratively adjusted *D*, starting from *D*^*′*^, until the simulated drug concentration for caspofungin, itraconazole, and amphotericin B at the edge of the ZOI matched the experimentally determined MFC value (Table III in Appendix B).

**TABLE III.** Minimum fungicidal concentration (MFC) and radius of zone of inhibition (RAD) of itraconazole, amphotericin B, and caspofungin for *C. auris* isolate 3.

| Isolate No. | Antifungal Drug | Antifungal Disk Mass ($\mu\text{g}$) | MFC ($\mu\text{g}/\text{ml}$) | RAD (mm) |
| --- | --- | --- | --- | --- |
| 3 | Itraconazole | 50 | 2 | 18 |
| 3 | Amphotericin B | 20 | 1 | 9 |
| 3 | Caspofungin | 5 | 1 | 14 |

Further details on determining *D* along with parameter values for modeling and simulation can be found in Appendix A. The code used for numerical simulation and data visualization was implemented in Python using the NumPy and Matplotlib libraries ([43]; see Data Availability).

## III. EXPERIMENTAL METHODS

### A. Strains, cell culture, and antifungal drugs

We acquired three clinical *C. auris* isolates from the Alberta Precision Laboratories -Public Health Laboratory. These isolates were stored in 25 % glycerol (Fisher Scientific, Canada, #BP229-1) at -80 ^*°*^C until needed. To revive the *C. auris* isolates, frozen stock was cultured on yeast extract-peptone-dextrose (YPD) agar plates and incubated at 35 ^*°*^C for 48 hours. The YPD agar medium was prepared by mixing 5 g of yeast extract (Sigma Aldrich, Canada, #8013-01-2), 10 g of bacto peptone (Gibco, USA, #211677), and 7.5 g of agar (Fisher Scientific, Canada, #9002-18-0) in 450 mL of distilled water. After autoclaving, 50 mL of a 2 % (w/v) dextrose (Fisher Scientific, Canada, #50-99-7) solution was added. Fresh subcultures were then prepared on Sabouraud dextrose agar (SDA; Millipore Sigma, USA, #1.05438.0500) plates and incubated at 35 ^*°*^C for 24 hours before performing BMDAs (to determine MIC and MFC) and DDAs (to determine tolerance).

Three antifungal drugs were selected for our study to evaluate dose-dependent antifungal tolerance in *C. auris*. Specifically, itraconazole (Sigma-Aldrich, Canada, #16657), amphotericin B (Sigma-Aldrich, Canada, #A9528), and caspofungin (Sigma-Aldrich, Canada, #179463-17-3). Stock solutions of itraconazole and amphotericin B were dissolved in DMSO (Fisher Scientific, Canada, #66-68-5), whereas caspofungin was dissolved in sterile water.

### B. Minimum inhibitory and fungicidal concentrations

1. *auris* isolate 3 was used to determine the *MIC*_50_ (MIC of a drug that inhibits the growth by 50 %, which we denote as MIC) and MFC of each antifungal drug used in our study. We used isolate 3 as we previously confirmed it to be susceptible to itraconazole, amphotericin B, and caspofungin [30]. To determine the concentration ranges at which tolerant *C. auris* colonies emerged (Sec. IV C), we used *C. auris* isolates 1 and 2, which we previously confirmed to exhibit tolerance to these three antifungal drugs [30].

The MIC of each antifungal drug was determined by performing a BMDA on the susceptible *C. auris* isolate 3 in accordance with the Clinical and Laboratory Standards Institute M27 guidelines [44]. Antifungal susceptibility testing was performed in 96-well Ubottom microplates (Thermo Fisher Scientific, Canada, #163320). The susceptible *C. auris* isolate was tested against itraconazole (0.03–16 µg/mL), amphotericin B (0.03–16 µg/mL), and caspofungin (0.015–8 µg/mL). *C. auris, Candida parapsilosis* (ATCC 22019), and *Issatchenkia orientalis* (ATCC 6258) were subcultured and incubated for 24 hours at 35 ^*°*^C prior to MIC testing; *C. parapsilosis* and *I. orientalis* were used as quality control strains. Each microwell was inoculated with 100 µL of a fungal suspension containing 2–5 *×* 10^3^ cells. Following inoculation, the microplates were incubated at 35 ^*°*^C and then the MICs were recorded after 24 and 48 hours. The MICs for the semi-solid DDAs were determined via the E-test strip method [45], using three antifungal strips placed on separate Muller-Hinton agar plates (MH: Sigma-Aldrich, Germany, #70191): itraconazole (Liofilchem, Italy, #LF92148), amphotericin B (Liofilchem, Italy, #LF92153), and caspofungin (Liofilchem, Italy, #LF92154).

The MFC was determined by first performing a BMDA in a 96-well microplate on the susceptible *C. auris* isolate 3. Each microplate well was inoculated with a *C. auris* inoculum prepared to match the MIC inoculum density, along with varying concentrations of an antifungal drug (Sec. III A). The microplates were incubated at 35 ^*°*^C for 24 hours. Following incubation, microwells that showed no visible growth and had concentrations equal to or higher than the MIC were selected. Next, samples from these wells were transferred to SDA plates and incubated at 35 ^*°*^C in absence of antifungal drugs. Finally, the MFC was determined by the lowest drug concentration at which no fungal colonies were observed by visual inspection of the SDA plates after 48 hours [46].

### C. Quantifying antifungal tolerance

Previously, we established the antifungal tolerance of our clinical *C. auris* isolates [30] used in this study via two approaches: FoG on Muller-Hinton (MH) (Sigma-Aldrich, Germany, #70191) agar medium and SMG in liquid medium [27, 32]. We use FoG in the present study to determine hill-type dose-response curves to the anti-fungal drugs (Sec. IV C). FoG was estimated using the *diskImageR* software [32] using DDA images acquired after 48 hours. FoG is defined as the fungal growth inside the ZOI normalized by the fungal growth outside the ZOI (i.e., drug-free lawn), and yields a dimensionless value between 0 and 1.

Our diffusion simulations provide the spatiotemporal drug concentrations *C*(*x, y, t*) for each antifungal drug used in this study. Assuming radial symmetry for the DDAs, we converted these 2D drug concentration profiles to a radial-temporal concentration profiles *C*(*r, t*). For each isolate-drug condition, we measured the radius of the closest (*R*_1_) and furthest (*R*_2_) tolerant colonies from the drug disk within the ZOI (Fig. 6 in Appendix B) at 48 hours. To determine the drug concentration ranges at which tolerant *C. auris* colonies emerged in each antifungal drug, these experimental radii were mapped to the corresponding simulated drug concentrations *C*_1_ = *C*(*R*_1_, 48 h) and *C*_2_ = *C*(*R*_2_, 48 h).

## IV. RESULTS AND DISCUSSION

### A. Diffusion constants for antifungal drugs

First, we calculated an initial diffusion constant (*D*^*′*^) for each antifungal drug using the Stokes-Einstein relation [Eq. (2)] and the radius [Eq. (3)]. The *D*^*′*^ for each drug differed due to differences in the radii [Eq. (2)] and molecular weights [Eq. (3)] of the drug molecules. These initial *D*^*′*^ estimates were about an order of magnitude smaller than the subsequent computationally determined *D* values (Sec. IV A; Table II in Appendix A). The *D*^*′*^ values were used as initial values to seed our numerical simulations, which we used to iteratively obtain more accurate *D* values for antifungal drug diffusion in our DDAs (Sec. II B).

Amphotericin B had the highest diffusion constant (*D*_*amp*_ = 9.54 *×* 10^*−*10^ m^2^ s^*−*1^) of all the antifungal drugs considered in this study, followed by itraconazole (*D*_*itr*_ = 6.01 *×* 10^*−*10^ m^2^ s^*−*1^), and then by caspofungin (*D*_*cas*_ = 3.00 *×* 10^*−*10^ m^2^ s^*−*1^). *D*_*cas*_ was the lowest of the three antifungal drugs, likely due to the larger molecular weight and size of capsofungin (Table II in Appendix A); this is supported by the fact that 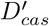 was the lowest of the Stokes-Einstein estimated values. *D*_*amp*_ was the highest of the antifungal drugs, despite the intermediate molecular weight, size, and 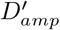 value of amphotericin The *D*_*itr*_ value was intermediate despite itraconazole having the lowest molecular weight and size, and the highest *D*^*′*^ value.

These results suggest that the computational *D* values are also determined by interactions between the physico-chemical properties and the polymer matrix of the agar medium [47]. These factors, as well as colony-drug interactions, were indirectly incorporated into our computationally determined *D* values, as these values were tuned to the experimental DDA data (Sec. II B). Therefore, we used the computational *D* values to simulate the spatiotemporal antifungal drug concentration profiles (Sec. IV B).

### B. Spatiotemporal antifungal drug concentration profiles

Next, we used the computationally determined *D* values (Sec. IV A) to simulate the spatiotemporal drug concentration profiles corresponding to our DDA experiments.

To quantify the diffusion of the antifungal drugs during the 48 hour DDAs, we plotted the concentration of each drug as a function of the RAD (Fig. 1). As in DDA experiments, drug diffusion started from different concentrations of itraconazole, amphotericin B, and caspofungin (Table II in Appendix A) from the center of the agar plate. The concentrations of all three drugs decreased sigmoidally to the edge of the ZOI (Fig. 1) until drug concentrations were approximately equal to MFC values after 48 hours (Table III in Appendix B).

**FIG. 1.**
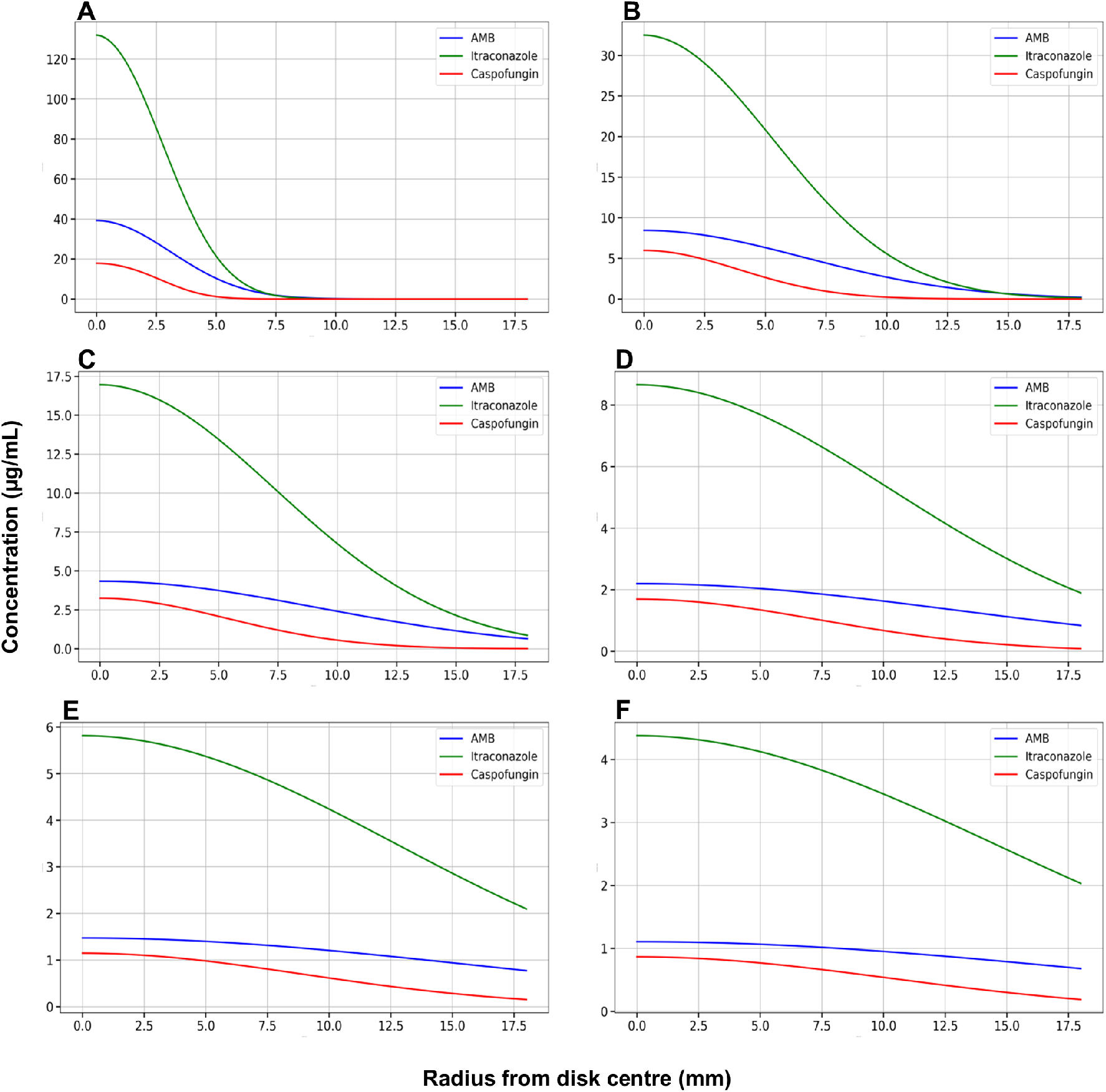
Radial-temporal antifungal drug concentration profiles from simulated disk diffusion assays. The concentration of itraconazole, caspofungin, and amphotericin B (AMB) as a function of radial distance from the drug disk after (A) 1 hour, (B) 6 hours, (C) 12 hours, (D) 24 hours, (E) 36 hours, and (F) 48 hours.

Throughout the simulations, the concentrations of amphotericin B, itraconazole, and caspofungin were highest near the drug disk and decreased as the drugs diffused toward the edge of the ZOI (Figs. 2-4). The radial concentrations changed more rapidly at early times in the simulations compared to later times (Fig. 1). Toward the end of the simulations, the spatial concentration profiles for itraconazole and caspofungin were nearly homogeneous (Fig. 1D-F), indicating that the spatial distribution of these drug molecules approached equilibrium in the DDAs.

**FIG. 2.**
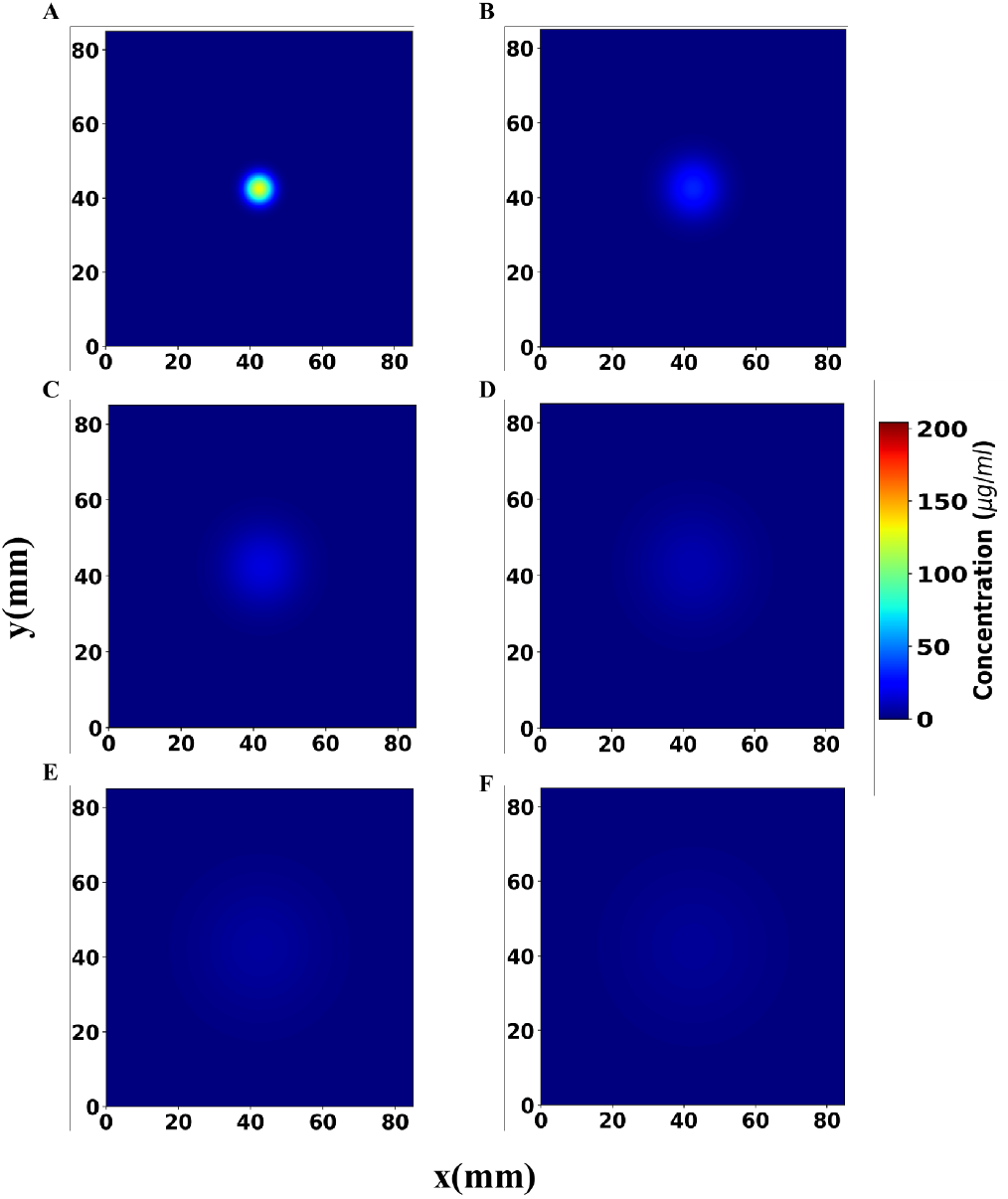
Spatiotemporal concentration profile of itraconazole from a simulated disk diffusion assay. Diffusion of itraconazole from a 50 *µ*g drug disk in x and y directions after (A) 1 hour, (B) 6 hours, (C) 12 hours, (D) 24 hours, (E) 36 hours, and (F) 48 hours. The diffusion constant was set to *D*_*itr*_ = 6.01 *×* 10^*−*10^ m^2^/s. The color bar denotes the concentration of itraconazole.

**FIG. 3.**
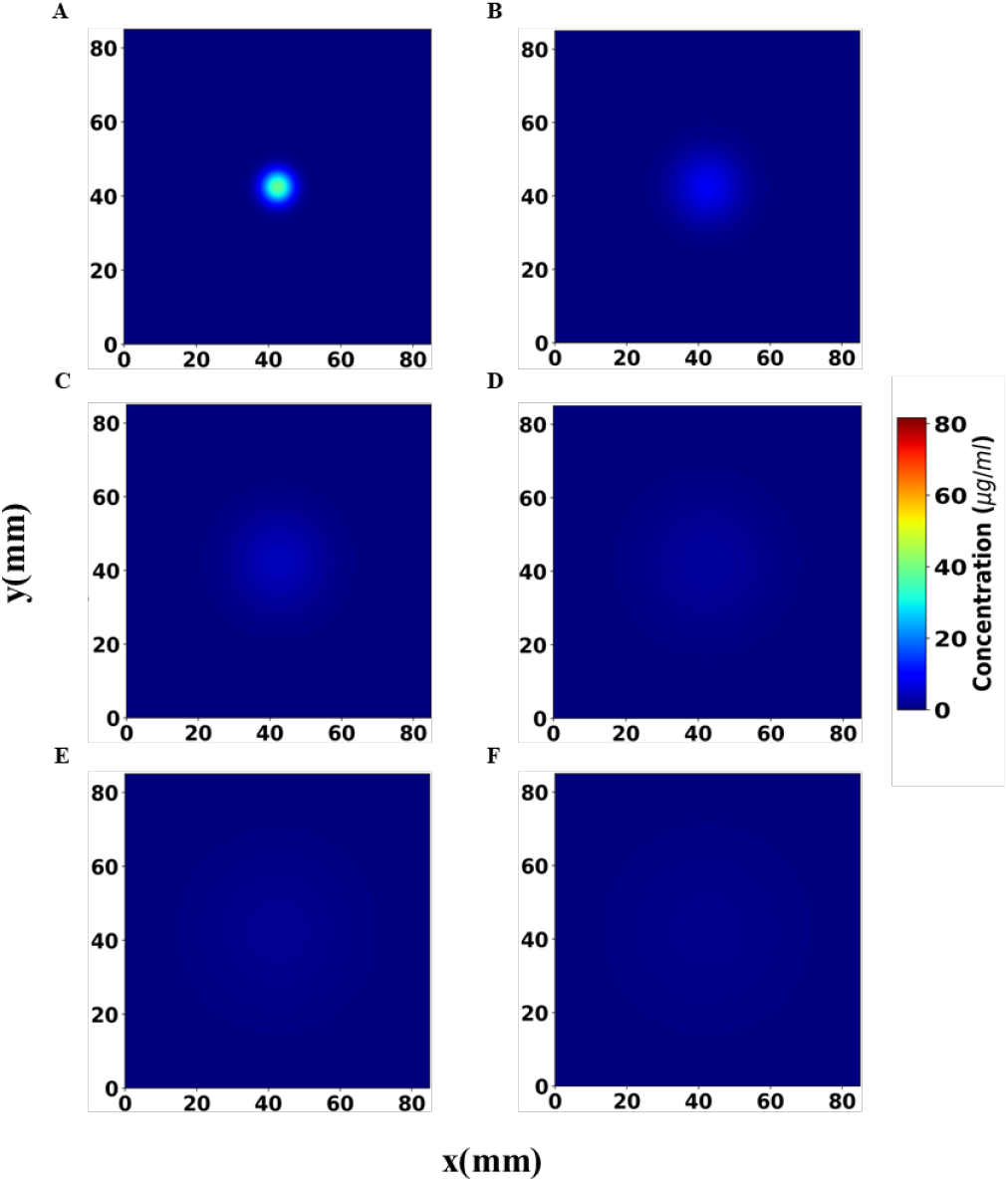
Spatiotemporal concentration profile of amphotericin B from a simulated disk diffusion assay. Diffusion of amphotericin B from a 20 *µ*g drug disk in the x and y directions after (A) 1 hour, (B) 6 hours, (C) 12 hours, (D) 24 hours, (E) 36 hours, and (F) 48 hours. The diffusion constant was set to *D*_*amp*_ = 9.54 *×* 10^*−*10^ m^2^/s. The color bar denotes the concentration of amphotericin B.

**FIG. 4.**
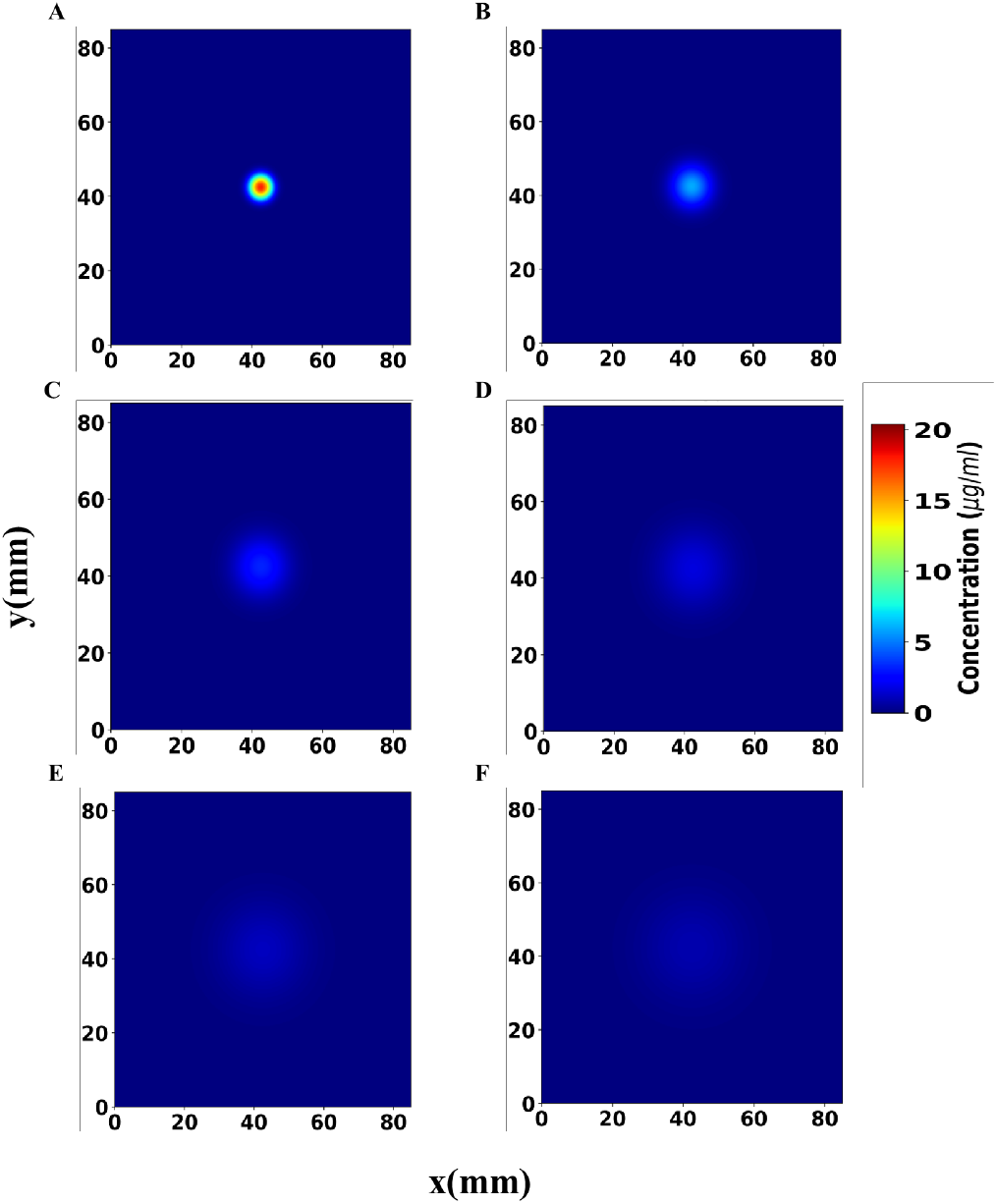
Spatiotemporal concentration profile of caspofungin from a simulated disk diffusion assay. Diffusion of caspofungin from a 5 *µ*g drug disk in x and y directions through agar after (A) 1 hour, (B) 6 hours, (C) 12 hours, (D) 24 hours, (E) 36 hours, and (F) 48 hours. The diffusion constant was set to *D*_*cas*_ = 3.00 *×* 10^*−*10^ m^2^/s. The color bar denotes the concentration of caspofungin.

The corresponding analytical solutions for radially symmetric scenarios are shown in Figure 7 in Appendix C). The numerical and analytical solutions both have radially decreasing concentration profiles that evolve monotonically over time (Fig. 1 and Fig. 7 in Appendix C, respectively). While the analytical solution (Eq. C3) diverges from the numerical solution at the beginning of the DDA (as *t →* 0; Fig. 1A and Fig. 7A in Appendix C), the analytical and numerical solutions are in quantitatively agreement at subsequent DDA time points (Fig. 1B-F and Fig. 7B-F in Appendix C).

The antifungal drug diffusion models provided the spatiotemporal drug concentration profiles that we used to quantify the dose-dependent nature of antifungal tolerance in *C. auris* (Sec. IV C).

### C. Dose-dependent antifungal tolerance in *C. auris*

Finally, we investigated how antifungal tolerance in *C. auris* depended on the drug concentration gradient within the ZOI of our DDAs. Using the experimentally calibrated concentration profiles of itraconazole, amphotericin B, and caspofungin (Sec. IV B), we identified the drug concentration ranges, or “tolerance windows”, within which tolerant *C. auris* colonies emerged; specifically, the drug concentration ranges corresponding to the tolerant-growth annulus in the ZOI. For each antifungal–isolate combination, tolerant colonies appeared within a distinct concentration range (Table I). This allowed us to map the experimental spatiotemporal emergence of tolerant colonies to the simulated drug-tolerance windows.

The percent change in *C*_1_ *− C*_2_ and *R*_1_ *− R*_2_ (Sec. III C; Table I) allowed us to quantitatively compare the emergence of tolerance in different isolate-drug conditions; these metrics indicate whether tolerance emerged over a relatively narrow or wide tolerance window and if the corresponding spatial distribution of tolerant colonies within the ZOI was relatively small or large, respectively. For the first *C. auris* isolate (ID1), a 35.3 % reduction in caspofungin concentration within the ZOI corresponded to a 56.3 % expansion in the radius of the tolerance annulus (Table I). Yet, for the second isolate (ID2), a 44.7 % decrease in caspofungin concentration resulted in a 127.3 % increase in the radius of the tolerance annulus. For third isolate (ID3), a 5.82 % reduction in amphotericin B concentration corresponded to a 22.2 % increase in the radius of the tolerance annulus. The fourth isolate (ID4) exhibited the most pronounced response, where a 32.7 % decrease in itraconazole concentration corresponded to a 154.6 % increase in the radius of the tolerance annulus. The tolerance windows for amphotericin B and caspofungin (*C*_1_ *− C*_2_ ranges in Table I) were below their corresponding MFC values (Table III in Appendix B). The MFC values were used as target drug concentrations at the edge of the ZOI after 48 hours to calibrate the simulations to obtain the more accurate *D* values (Sec. II B). This discrepancy occurred due to the limited logarithmic sampling range (optimized to balance computational cost and accuracy) used to computationally determine the *D* values (Appendix A), which resulted in simulated drug concentrations below MFC at the edge of the ZOI after 48 hours for these two antifungal drugs (Fig. 1).

For each isolate-drug condition, *diskImageR* [32] analysis of the DDA images yielded the FoG values, while the calibrated diffusion model provided the corresponding antifungal drug concentrations within the ZOI. Mapping the experimental tolerant-colony annulus (*R*_1_-*R*_2_) onto the simulated concentration profile generated a bounded tolerance window (*C*_1_ - *C*_2_) within which the tolerant colonies emerged (Table I). To provide a FoG-drug concentration analysis of these results, we constructed Hill-type dose response representations [26] by pairing the experimentally derived FoG values with the corresponding simulation-derived drug concentrations (Fig. 5). These plots highlight that tolerant growth occurs above MIC over a finite antifungal concentration range, and that the width and position of tolerance windows depend on the isolate and the antifungal drug class. For itraconazole and amphotericin B, some of the *RAD*_20_, *RAD*_50_, *RAD*_80_, and values were on the edge or outside of the tolerance windows. This resulted from *diskImageR* incorporating all colonies that emerged within the ZOI at up to the 48 hour time points in the FoG analysis (which were used to determine the FoG and *RAD*_20_, *RAD*_50_, and *RAD*_80_ values in Fig. 5), whereas we incorporated colonies within the ZOI after 24 hours (which were used to determine the tolerance windows in Fig. 5).

**FIG. 5.**
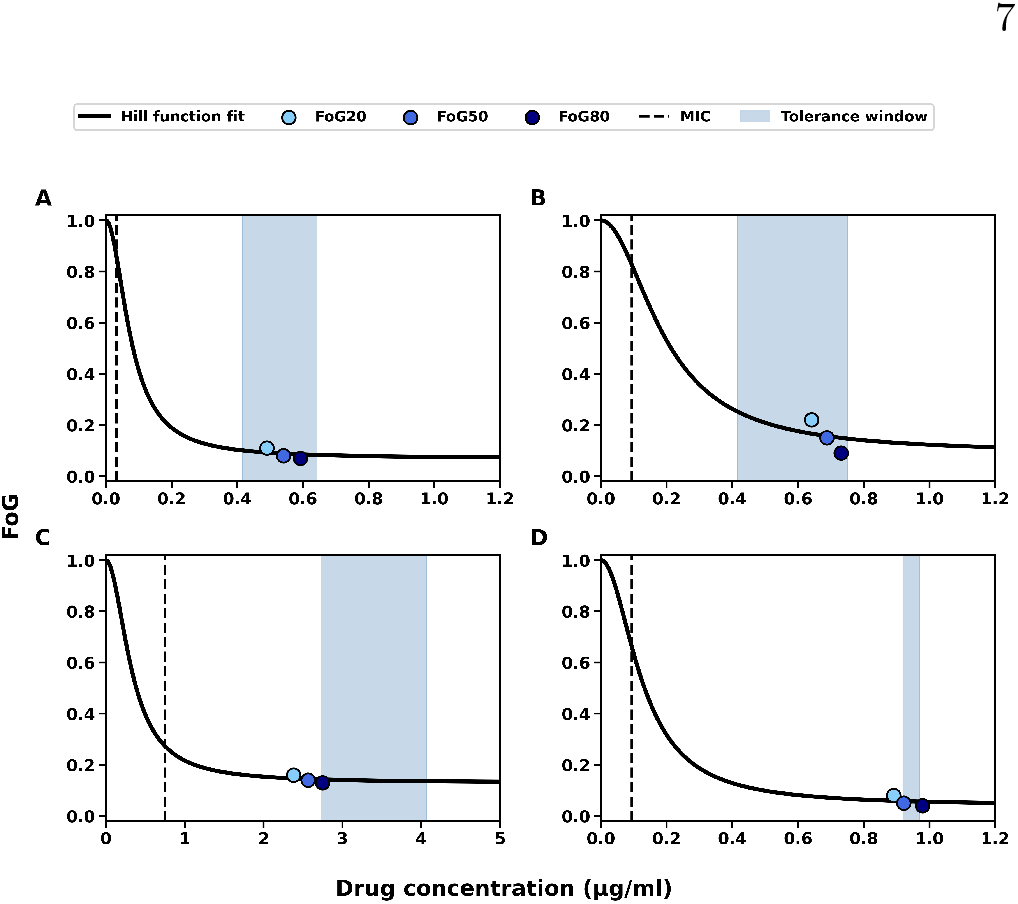
Hill-type dose–response representation of antifungal tolerance in *Candida auris*. Panels (A)–(D) show the relationship between the fraction of growth inside the zone of inhibition (FoG) and antifungal drug concentration for different isolate–drug combinations: (A) isolate 1 with caspofungin (MIC = 0.032 *µ*g/ml), (B) isolate 2 with caspofungin (MIC = 0.094 *µ*g/ml), (C) isolate 1 with itraconazole (MIC = 0.75 *µ*g/ml), and (D) isolate 1 with amphotericin B (MIC = 0.094 *µ*g/ml). The dashed vertical line in each panel indicates the MIC, and the blue shaded region represents the simulated drug tolerance window for the colonies appeared at 48 hours. In each panel, the three data points correspond to the radius of the zone of inhibition (RAD) where 80 %, 50 %, and 20 % of growth reduction occur: RAD_80_ (highest drug concentration, lowest FoG; dark blue dot), RAD_50_ (intermediate drug concentration and FoG; medium blue dot), and RAD_20_ (lowest drug concentration, highest FoG; light blue dot). Solid lines are the Hill function fit to the FoG data. See Appendix A for details of the model and fitting procedure.

**FIG. 6.**
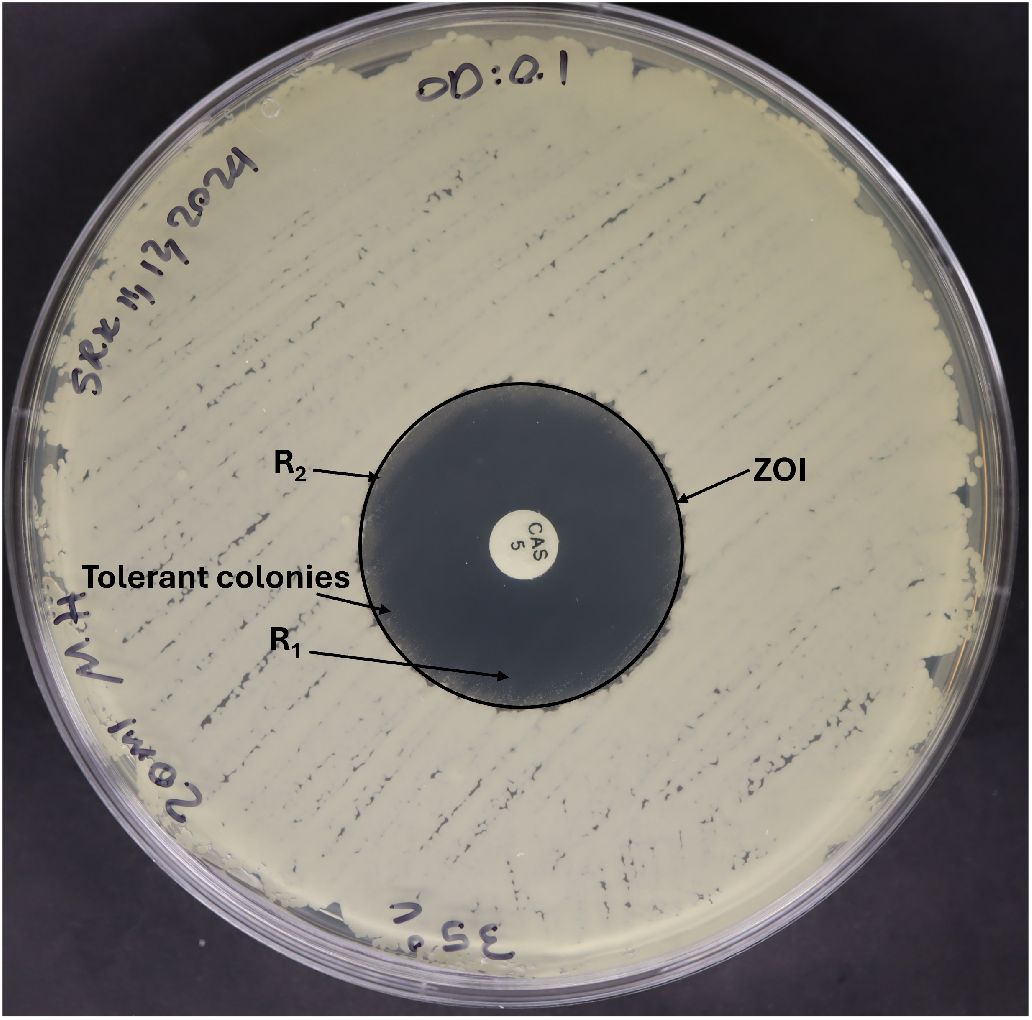
Tolerant *C. auris* isolate 1 colonies in the zone of inhibition of a 5 *µ*g caspofungin (here denoted CAS) disk diffusion assay after 48 hours.

**FIG. 7.**
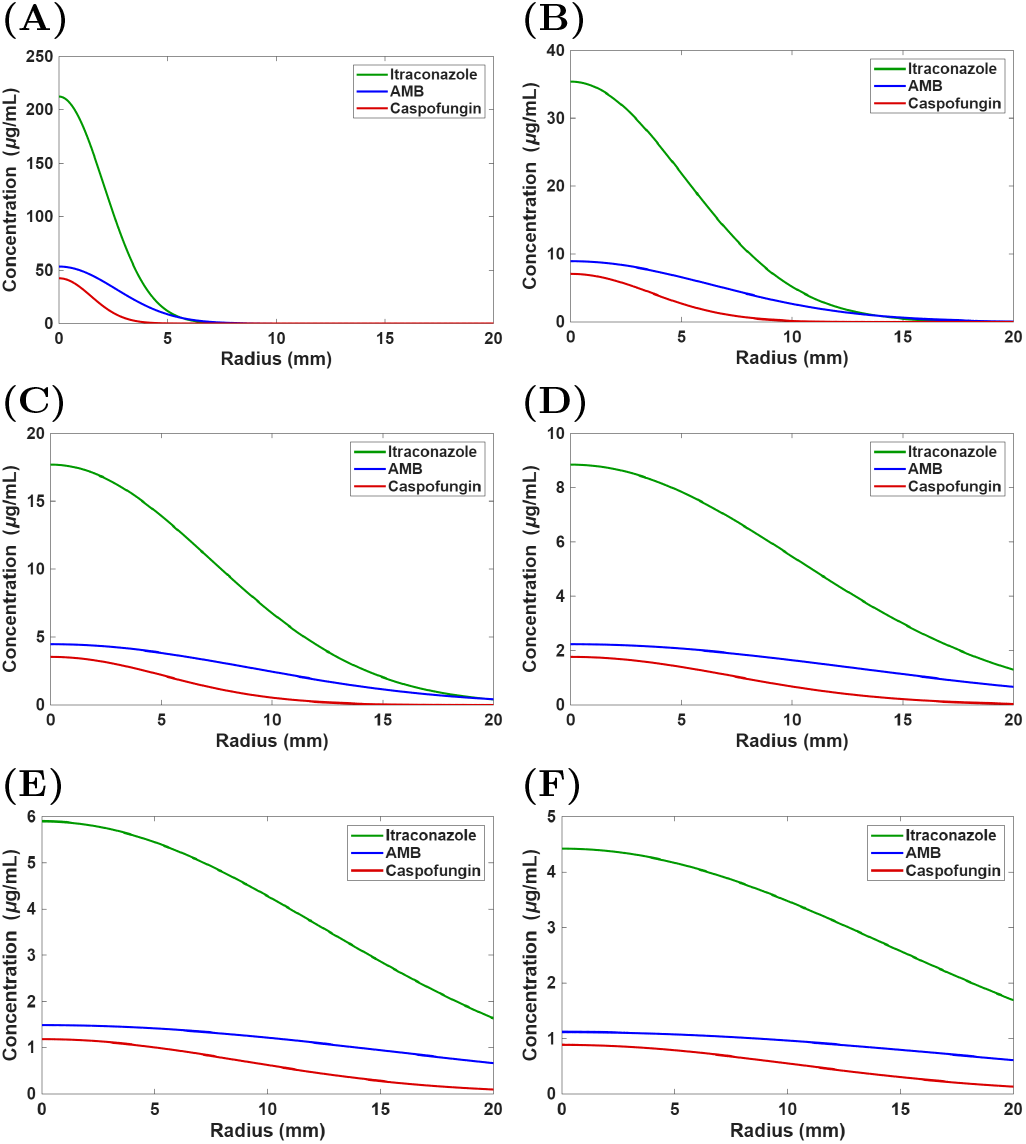
Analytical solutions for diffusion of caspofungin, itraconazole, and amphotericin B (AMB) after (A) 1 hour, (B) 6 hours, (C) 12 hours, (D) 24 hours, (E) 36 hours, and (F) 48 hours.

Overall, these results demonstrate that antifungal tolerance in *C. auris* is a strain-dependent, drug-dependent, and dose-dependent phenomenon.

## V. CONCLUSION

We discovered that antifungal tolerance is a strain-, drug-, and dose-dependent phenomenon in a multidrug-resistant human fungal pathogen. In our experimental DDAs, tolerant *C. auris* colonies grew near the outer edge of the ZOI during exposure to all three major classes of antifungal drugs. Using our computational-experimental approach, we determined that these tolerant colonies emerged within diffusion-mediated drug concentration windows that differed between clinical isolates and antifungal drug classes. This contrasts with “classical tolerance” where tolerant colonies emerge homogeneously throughout the ZOI [27, 30].

We combined established methods from physics and microbiology to quantify the emergence of antifungal tolerance on drug concentration. The Stokes-Einstein relation was used to calculate an initial value of the diffusion constants for the antifungal drugs used in our study. Although the Stokes-Einstein relation accounts for temperature, viscosity, and molecular radius, our drug diffusion simulations using these initial diffusion constants did not agree with our experimental DDA data. To improve the accuracy of our numerical simulations, we recalibrated the diffusion constants to satisfy a simulation boundary condition; the value of the diffusion constant was iteratively changed until the drug concentration at the edge of the simulated ZOI after 48 hours was equal to the experimentally determined MFC. By integrating experimental DDA and MFC data into our drug diffusion modeling, we indirectly incorporated unmeasured physical and biochemical factors, such as solubility, pH, and temperature, which would be resource intensive to determine experimentally. However, it remains to be elucidated how these physical and biochemical factors impact tolerance. We focused on numerical simulations, as opposed to analytical solutions (Appendix C), to enable future computational research on the effects of non-uniformly distributed colonies and more realistic 3D drug diffusion on antifungal tolerance.

For the clinical *C. auris* isolates that we investigated, higher concentrations of antifungal drugs inhibited growth, while lower concentrations allowed the survival and proliferation of tolerant colonies in our DDAs. We quantified the drug concentration ranges, or “tolerance windows”, for the emergence of tolerant colonies using our computational-experimental approach. We found an intermediate tolerance window for the azole itraconazole, a narrow tolerance window for the polyene amphotericin B, and an intermediate to wide, strain-dependent tolerance window for the echinocandin caspofungin. These differences in these tolerance windows may be due to the unique mechanism of action of each drug class [24], however, this remains to be elucidated in larger-scale molecular studies.

Similar to the transition from non-genetic to genetic drug resistance [48, 49], exposure to tolerance windows may enrich infections for tolerant subpopulations that evolve to be genetically drug resistant. Quantitatively understanding how drug diffusion influences antifungal tolerance *in vivo* may improve clinical outcomes by informing dosing and drug combination strategies [26, 27] to avoid tolerance windows before irreversible and life-threatening antifungal resistance evolves.

## ACKNOWLEDGMENTS

We thank Dr. Tanis Dingle at the Alberta Health Services — Public Health Laboratory for providing the clinical *C. auris* isolates. We also thank the Molecular Biology Services Unit at the University of Alberta for their help with genetic sequencing. We acknowledge Prof. Judith Berman, Nora Kawar, and Dr. Akila Bandara for helpful discussions, Danial Papi for assistance with Python programming, Harold Flohr for assistance with error analysis, and Joshua Guthrie for assistance with LaTeX formatting. D.A.C. was supported financially by a Government of Canada’s New Frontiers in Research Fund — Exploration grant (NFRFE-2019-01208) and an Audrey and Randy Lomnes Early Career Endowment Award. C.M.G. was funded by the DOST-SEI-UAlberta S&T Graduate Scholarship Program.

## AUTHOR CONTRIBUTIONS

D.A.C. conceptualized the study. D.A.C. and S.R.K. designed the study. S.R.K. performed the experiments and numerical simulations. C.M.G. performed the analytical modeling. S.R.K., D.A.C., and C.M.G. analyzed the results. D.A.C., S.R.K., and C.M.G. wrote the manuscript. D.A.C. acquired the funding and supervised the study.

## DATA AVAILABILITY

The Python code used to perform the numerical simulations and the experimental data supporting the findings of this study are available at: https://data.mendeley.com/preview/m9t5f3ryvz?a=a1e8e284-00d4-4f49-8b58-c9eaf0b65385.

## Appendix A Parameters & Modeling

The parameters for the modeling and simulations conducted in our study are listed in Table II.

For each antifungal drug, Eq. (2) was solved iteratively using trial values of the diffusion coefficient, *D*^*′′*^. *D*^*′′*^ values were sampled logarithmically over a predefined range, starting from a seed value, *D*^*′*^ (Sec. II A). For each value of *D*^*′′*^, the radial concentration profile *c*(*r, t*; *D*^*′′*^) was computed at *t* = 48 h. The average antifungal drug concentration at the experimentally measured RAD was then evaluated and compared with the corresponding MFC. The difference in these values was then determined:

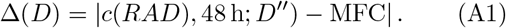

The *D*^*′′*^ that minimized Eq. (A1) was selected as the calibrated *D* value for each antifungal drug (Table II). This procedure was performed independently for itraconazole, amphotericin B, and caspofungin.

The Hill function fit in Figure 5 was determined using the following equation:

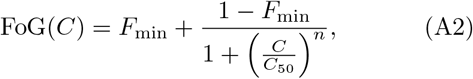

where *C* is the antifungal drug concentration. The parameters of the Hill function were estimated separately for each isolate–drug combination shown in Fig. 5. The estimated half-maximal concentration (*C*_50_) was 0.077 *µ*g/ml for caspofungin (isolate 1), 0.194 *µ*g/ml for caspofungin (isolate 2), 0.333 *µ*g/ml for itraconazole (isolate 1), and 0.127 *µ*g/ml for amphotericin B (isolate 1). The corresponding minimum FoG (*F*_min_) values, which were obtained from the lowest FoG value in each dataset, were 0.07, 0.09, 0.13, and 0.04, respectively. The Hill coefficient was set to *n* = 2 for all cases.

## Appendix B Supporting Experimental Information

Our diffusion model assumed a two-dimensional radial spread of antifungal drugs on the agar surface (Sec. II A). To examine whether the depth of the agar medium affects this assumption, we performed DDAs using different volumes of MH agar. Among the three antifungals tested, only amphotericin B had a statistically significant change in RAD across agar volumes [*F* (3, 8) = 12.92, *p* = 0.002]. This statistical effect was primarily driven by the thin agar layer (10 ml) condition, which resulted in a larger RAD compared to the thicker agar layer (15, 25, and 30 ml) conditions. In contrast, the RADs across agar volumes for caspofungin and itraconazole did not show significant differences (Table IV). These results support the use of a two-dimensional diffusion model in this study.

**TABLE IV.** Radius of the zone of inhibition (RAD) for itraconazole, caspofungin, and amphotericin B disk diffusion assays for *C. auris* isolate 1 using different volumes of Muller-Hinton agar. A one-way ANOVA showed no significant effect of volume on RAD for itraconazole (*p* = 0.370) and caspofungin (*p* = 0.361); however, a significant effect was found for amphotericin B (*p* = 0.002). Tukey’s HSD test revealed that the 10 ml agar condition was significantly different from the 15 ml (*p* = 0.009), 25 ml (*p* = 0.011), and 30 ml (*p* = 0.002) conditions, while differences between the 15 ml versus 25 ml (*p* = 0.999), 15 ml versus 30 ml (*p* = 0.518), and 25 ml versus 30 ml (*p* = 0.453) conditions were not statistically significant for all antifungal drugs.

| Agar Volume (ml) | Caspofungin (RAD Mean ± SD, mm) | Itraconazole (RAD Mean ± SD, mm) | Amphotericin B (RAD Mean ± SD, mm) |
| --- | --- | --- | --- |
| 10 | 16.09 ± 0.38 | 19.85 ± 0.12 | 14.57 ± 0.31 |
| 15 | 15.44 ± 0.49 | 21.20 ± 1.66 | 13.15 ± 0.39 |
| 25 | 15.51 ± 0.55 | 20.21 ± 0.52 | 13.19 ± 0.44 |
| 30 | 15.67 ± 0.37 | 20.19 ± 0.61 | 12.70 ± 0.40 |

Viscosity measurements were carried out using a Modular Compact Rheometer (Anton Paar GmbH, Graz, Austria, MCR 102) in the Department of Chemical and Materials Engineering at University of Alberta. To measure the viscosity of agar medium, a shear rate of 1000 s^*−*1^ was applied and measured 10 times per minute at 35 °C.

## Appendix C Analytical Solution for Drug Diffusion

As presented in Sec. II A, antifungal drug diffusion in a DDA can be modeled using Fick’s second law in two-dimensions. Transforming Eq. (1) to polar coordinates, we obtain:

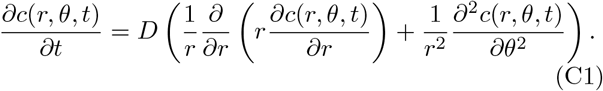

Since the drug disk is circular and the agar medium in the Petri dish is assumed to be homogeneous, drug diffusion here is radially symmetric. By circular symmetry (*∂c/∂θ* = 0), Eq. (C1) reduces to a 1D radial diffusion equation [52]:

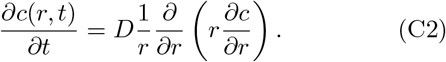

For an initially localized drug source (i.e., an antifungal drug in a paper disk approximated as a point source of total mass *M*) in an infinite two-dimensional medium is:

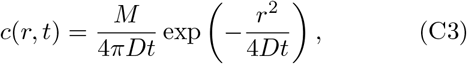

where *r* is the radial distance from the center of the drug disk. *M*_*itr*_ = 50 *µ*g, *M*_*cas*_ = 20 *µ*g, and *M*_*amp*_ = 5 *µ*g. The total mass *M* in the drug disk can be computed as: *M*_*i*_ = *C*_*i*_ *× V*_*i*_, where *∈ {i itr, amp, cas}* . *D* values were obtained from Table II. The solutions to Eq. (C3) for each antifungal drug are shown in Fig. (7).

Overall, the analytical solutions (Fig. 7) for the antifungal drugs considered in this study are in agreement with the numerical solutions (see Sec. IV B; Fig. 1).

